# Taking rapid and intermittent cocaine infusions enhances both incentive motivation for the drug and cocaine-induced gene regulation in corticostriatal regions

**DOI:** 10.1101/2020.04.22.055715

**Authors:** Ellie-Anna Minogianis, Anne-Noël Samaha

## Abstract

A goal in addiction research is to distinguish forms of neuroplasticity that are involved in the transition to addiction from those involved in mere drug taking. Animal models of drug self-administration are essential in this context. Here, we compared in male rats two cocaine self-administration procedures that differ in the extent to which they evoke addiction-like behaviours. We measured both incentive motivation for cocaine using progressive ratio procedures, and cocaine-induced c-*fos* mRNA expression, a marker of neuronal activity. Rats self-administered intravenous cocaine (0.25 mg/kg/infusion) for seven daily 6-hour sessions. One group had intermittent access (IntA; 6 minutes ON, 26 minutes OFF x 12) to rapid infusions (delivered over 5 seconds). This models the temporal kinetics of human cocaine use and produces robust addiction-like behaviour. The other group had Long access (LgA) to slower infusions (90 seconds). This produces high levels of intake without promoting robust addiction-like behaviour. LgA-90s rats took twice as much cocaine as IntA-5s rats did, but IntA-5s rats showed greater incentive motivation for the drug. Following a final self-administration session, we quantified c-*fos* mRNA expression in corticostriatal regions. Compared to LgA-90s rats, IntA-5s rats had more cocaine-induced c-*fos* mRNA in the orbitofrontal and prelimbic cortices and the caudate-putamen. Thus, a cocaine self-administration procedure (intermittent intake of rapid infusions) that promotes increased incentive motivation for the drug also enhances cocaine-induced gene regulation in corticostriatal regions. This suggests that increased drug-induced recruitment of these regions could contribute to the neural and behavioural plasticity underlying the transition to addiction.

## INTRODUCTION

Drug addiction is a chronic and relapsing disorder defined by recurring drug-seeking and drug-taking in spite of adverse consequences (APA, 2013). Many people use drugs, but only a subset develop addiction (Anthony et al., 1994; Gawin, 1991; Shaffer and Eber, 2002). A challenge is to distinguish the changes in brain, psychological function and behaviour that underlie the development of addiction from those that result from mere drug taking. To this end, drug self-administration procedures have been developed, whereby laboratory animals not only voluntarily take drug, but also develop behavioural features relevant to addiction, including enhanced responding for drug when cost is high (e.g., under a progressive ratio schedule of reinforcement; PR), continued responding for drug under extinction conditions and/or in spite of adverse consequences (e.g., footshock), and augmented relapse-like behaviour.

Self-administration procedures have manipulated how much drug gets to the brain, how fast, and how often (reviewed inAllain et al., 2015). For instance, taking rapid, rather than more sustained i.v. cocaine infusions promotes addiction-relevant patterns of drug use. Rats taking faster i.v. infusions of cocaine [injected over 5 versus 90 seconds (s)] consume more drug (Allain et al., 2018; Bouayad-Gervais et al., 2014; Minogianis et al., 2013; Wakabayashi et al., 2010), more readily develop psychomotor sensitization (Allain et al., 2017), respond more for cocaine under a PR schedule (Allain et al., 2017; Bouayad-Gervais et al., 2014; Liu et al., 2005; Minogianis et al., 2013), and show enhanced relapse-like behaviour following abstinence (Gueye et al., 2019; Wakabayashi et al., 2010).

Rats given intermittent (IntA) rather than continuous (long access, or LgA) cocaine access during each self-administration session are also more likely to develop an addiction-like phenotype (reviewed in Allain et al., 2015; Kawa et al., 2019; Zimmer et al., 2012). During an IntA session (4-6 hours), drug-available periods are separated by longer, no drug-available periods (Zimmer et al., 2011). This produces the peaks and troughs in brain cocaine concentrations that are thought to model the temporal pattern of human cocaine use (Beveridge et al., 2012; Zimmer et al., 2011). LgA rats take much more drug, but IntA rats show more incentive motivation for cocaine, as measured either by behavioural economic indicators or responding for the drug under PR (Algallal et al., 2019; Allain et al., 2018; Kawa et al., 2016; Zimmer et al., 2012). IntA rats also persist in responding for cocaine in spite of an adverse consequence (i.e., mild footshock), they seek the drug when it is not available, and they show stronger cue-induced relapse behaviour than generally seen in LgA-rats (Gueye et al., 2019; Kawa et al., 2016; Nicolas et al., 2019; Singer et al., 2018).

To produce addiction, drugs are thought to change synapse organization and evoke long-lasting forms of neurobehavioural plasticity. Drug-induced changes in immediate early gene expression is thought to be an initial step in this plasticity (Hyman et al., 2006; Nestler, 2001). For instance, repeated exposure to drugs alters synaptic organization in regions of the brain involved in incentive motivation and reward, such as the striatum, and regions of the brain involved in judgement and inhibitory control of behaviour, such as the frontal cortex (Jentsch and Taylor, 1999; Robinson and Berridge, 2003). Some of this drug-induced plasticity produces sensitization to the incentive motivational effects of drugs and augments drug wanting, thereby promoting the transition to addiction (Robinson and Berridge, 1993; 2003).

Thus, animals are more likely to develop addiction-like behaviours if they take rapid versus slower cocaine injections and if they take the drug intermittently rather than continuously during each self-administration bout. Here we leveraged these pharmacokinetic principles to compare incentive motivation for cocaine and cocaine-induced gene regulation following a pattern of cocaine use that readily produces an addiction-like phenotype (intermittent intake of rapid cocaine infusions), versus a pattern of use that does not (continuous intake of slower cocaine infusions).

## MATERIALS AND METHODS

### Animals

Forty male Wistar rats (Charles River Laboratories, St-Constant, QC) weighing between 225-250 g upon arrival were housed individually in a climate-controlled colony room maintained on a reverse 12 h/12 h light/dark cycle (lights off at 8:00 am). Experiments were conducted during the dark phase of the rats’ circadian cycle. Food and water were available *ad libitum*, until the day after surgery, from which point on, animals received 5 food pellets at the end of the day (25 g). Mild food restriction produces healthier rats compared to ad libitum feeding, which promotes excessive fat deposition and obesity (Martin et al., 2010; Rowland, 2007). The animal care committee of the Université de Montréal approved all procedures (Protocol #14-149).

### Drugs

Cocaine hydrochloride (Medisca Pharmaceutique Inc, St-Laurent, QC) was dissolved in 0.9% physiological saline (Medical Mart, Mississauga, ON) and filtered with Corning bottle-top filters (0.22 μm PES membrane; Fisher Scientific, Whitby, ON).

### Surgery and operant cages

**Figure 1** depicts the timeline of experimental events. After 1 week of habituation to the vivarium, a catheter was implanted into the jugular vein of each rat (Samaha et al., 2011; Weeks, 1962). To avoid blood clots in the catheters, they were flushed on alternate days with either 0.1 ml physiological saline or saline containing 0.2 mg/ml Heparin (Sigma-Aldrich, Oakville, ON) and 2mg/ml enroflaxin (Baytril®, CDMV, St-Hyacinthe, QC). On the day before progressive ratio testing, and on the day when brains were extracted (see ‘*Final cocaine self-administration session and brain extraction*’), catheter patency was verified by i.v. infusion of a sodium thiopental/sterile water solution (0.2 ml of a 20 mg/ml solution; CDMV, St-Hyacinthe, QC). All animals became ataxic within 5 s of the infusion and were included in data analyses. At least one week after surgery, the rats were trained to press a lever to obtain i.v. cocaine (0.25 mg/kg/infusion) in standard operant chambers (Med Associates, St-Albans, VT). The chambers were in a room separate from the rats’ housing room. Each chamber was placed within a light and sound attenuating cabinet equipped with a ventilation fan that also masked external noise. Infusion pumps with 3.33-RPM motors were used to deliver cocaine over 5 or 10 s at a rate of 29.68 μl/s, while 0.1-RPM motors delivered cocaine over 90 s at a rate of 0.803 μl/s. Each chamber contained two retractable levers and 4 photobeam sensors (Med Associates, St-Albans, VT) to measure locomotor activity. A computer running Med Associates Med-PC version IV software (Med Associates, St-Albans, VT) controlled parameters in the chambers and collected data.

**Figure 1.**
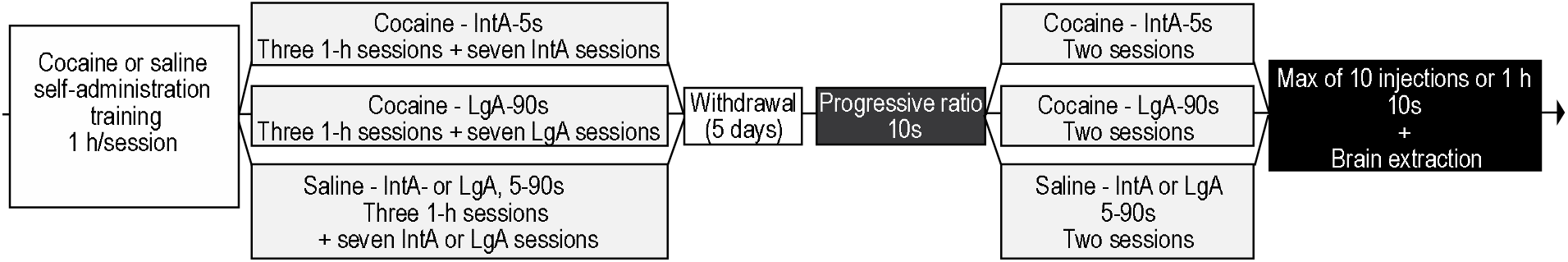
Timeline of experimental events. Rats were trained to self-administer cocaine (0.25 mg/kg/infusion) or saline delivered intravenously over 5 or 90 s seconds during 1-hour sessions, under a fixed ratio schedule of reinforcement. Following three more 1-hour sessions at their respective infusion speed, rats were further divided into three groups; IntA-5s rats were given intermittent access to rapid cocaine infusions (5 s), LgA-90s rats were given continuous access to slower cocaine infusions (90 s), and saline control rats self-administered saline under either IntA or LgA conditions. Five days following the cessation of self-administration, motivation for cocaine (0.063-0.25 mg/kg/infusion) or saline (saline control rats) was assessed using a progressive ratio schedule of reinforcement (PR). During PR sessions, all infusions were delivered over 10 s, such that testing conditions were held constant across groups. After PR testing, rats were given 2 more IntA or LgA sessions. On the following day, rats received a final cocaine or saline (saline control rats) self-administration session. This session ended after rats had taken 10 infusions or after 1 hour. Brains were then extracted. *h*, hour. *IntA*, intermittent access. *LgA*, long access. *s*, seconds.

### Self-administration training

After 7 days of recovery from surgery, rats were trained to self-administer cocaine (0.25 mg/kg/infusion) delivered over 5 s, during daily 1-h sessions. Sessions began with the illumination of the house light and insertion of both levers. Projected IntA-5s rats were trained to lever-press for cocaine under a fixed ratio (FR) 3 schedule of reinforcement. During each 5-s infusion and for 20 s thereafter, both levers were retracted and the light located above the active lever was turned on, as in (Allain et al., 2018). Projected LgA-90s rats were trained to lever-press for cocaine under FR2, with the timeout period gradually increasing from 20 s to 85 s, so that these rats would get used to waiting 90 s between injections, as in (Minogianis et al., 2013). We have previously shown that under these training conditions, LgA-90s rats acquire and then maintain reliable cocaine self-administration, but they do not develop strong incentive motivation to take the drug (Minogianis et al., 2013). We used FR3 instead of FR2 in the projected IntA-5s rats because in our experience, this increases discrimination between the active and inactive levers in these rats (Allain et al., 2018). Of note, when tested under a PR schedule, IntA rats show more incentive motivation for cocaine than LgA rats do, even when both groups had previously self-administered the drug under a common schedule of reinforcement (i.e., FR3; Allain et al., 2018). Pressing on the inactive lever had no programmed consequences. Rats were considered to have acquired reliable cocaine self-administration behaviour if, for 2 consecutive sessions they *i*) took ≥ 5 infusions/session, *ii*) at regular time intervals during the session (as determined by looking at cumulative response records), and *iii*) if they pressed twice as much on the active versus inactive lever. In parallel to the cocaine-taking rats, a third group of animals from the same cohort self-administered i.v. saline injections. Following acquisition of cocaine self-administration behaviour, all rats were given 3 additional 1-h sessions (1 session/day). During these sessions, the projected IntA-5s rats self-administered cocaine (0.25 mg/kg/infusion, delivered over 5 s) under FR3, with no timeout period. The projected LgA-90s rats self-administered cocaine under FR2, but each infusion was now delivered over 90 s, with no timeout period. The control rats continued to self-administer saline. During each infusion, the light above the active lever was turned on. After these 3 sessions, all rats were given 7 daily, 6-h sessions. During these sessions, the projected IntA-5s rats now self-administered 5-s cocaine infusions under IntA. The projected LgA-90s rats now self-administered 90-s cocaine infusions under LgA. One half of the control rats self-administered saline under IntA, and the other half self-administered saline under LgA, where infusions were delivered over 5-90s.

### Self-administration under intermittent access (IntA) conditions

Each IntA session had twelve, 6-min drug periods separated by 26-min no-drug periods during which levers were retracted and cocaine was not available. During each 6-min drug period, the animals could self-administer a maximum of 2 infusions (0.25 mg/kg/infusion; under FR3). Under these conditions, IntA rats show more incentive motivation for cocaine than LgA rats (Allain et al., 2018). Once the two infusions were self-administered or the 6-min drug period had elapsed, a 26-min no-drug period was initiated. Thus, the animals could take a maximum of 24 infusions per IntA-session. The last 6-min drug period was followed by a 2-min no-drug period such that the session lasted no more than 6 h.

### Self-administration under long access (LgA) conditions

During each self-administration session, cocaine was available continuously (0.25 mg/kg/infusion; under FR2), except during each 90-s infusion, where further lever presses did not produce additional infusions.

### Modeling brain cocaine concentrations

We estimated brain cocaine concentrations (μM) using self-administration data from the last (7^th^) IntA-5s and LgA-90s session, using a representative rat from each group, as in previous studies (Allain et al., 2018; Zimmer et al., 2011; Zimmer et al., 2012). We used a mathematical model developed and validated in male rats (Pan et al., 1991). The Python script used to model brain cocaine concentrations was kindly provided by Dr. David C. S. Roberts.

### Cocaine self-administration under a progressive ratio schedule of reinforcement (PR)

Five days after the last IntA or LgA session, we assessed incentive motivation for cocaine by allowing the rats to self-administer the drug (0.063-0.25, in descending order, 1-2 sessions/dose) under PR. Control rats were tested in parallel and self-administered saline under a PR schedule. During each PR session, the number of lever presses required to obtain each successive infusion increased exponentially, according to the formula (5 e^(injection number × 0.2)^ − 5), as described by (Richardson and Roberts, 1996). Cocaine (or saline) infusions were delivered over 10 s. Thus, dose and injection speed were held constant across groups, such that any group differences in responding for cocaine would be due to drug-taking history. Each PR session ended when an hour had elapsed since the last infusion or after 5 hours. The total number of active lever-presses was used as a measure of motivation for cocaine. The day after the last PR session, the rats received 2 final IntA-5s or LgA-90s sessions. Control rats received 2 IntA or LgA saline self-administration sessions.

### Final cocaine self-administration session and brain extraction

One day after the last IntA-5s or LgA-90s session, all rats received a final self-administration session before brains were extracted for *in situ* hybridization of c-*fos* mRNA. During this session, cocaine (or saline) was available under a FR2 schedule of reinforcement, and each infusion was administered over 10 s. To avoid the potentially confounding influence of acute differences in drug intake on c-*fos* mRNA density, all rats were allowed to take a maximum of 10 infusions during this final session. The session ended once the 10 infusions were taken or after 1 hour. Rats were sacrificed 45 minutes after the end of the session, because *c-fos* mRNA levels peak 45-60 min after i.v. cocaine administration (Ennulat et al., 1994; Moratalla et al., 1993). The rats received an i.v. infusion of a sodium thiopental/sterile water solution, they were then decapitated while anesthetized. The brains were extracted, plunged into cold isopentane (−50°C) and stored at −80°C.

### In situ *hybridization*

A subset of representative rats from each group was used for *in situ* hybridization (IntA-5s and LgA-90s, n = 6/group; Sal-Ctrl, n = 10), to keep the hybridization procedure manageable. Self-administration data in this subset was comparable to that of the main groups (see Results). *C-fo*s mRNA expression was labelled on 12-μm-thick coronal brain sections using a [^35^S]-UTP-labelled riboprobe complementary to *c-fos*, as in (Bédard et al., 2011). The complementary RNA probe for *c-fos* mRNA was derived from 1.8 kb *EcoRI* fragment of a full-length rat *c-fos* cDNA. It was subcloned into pBluescript SK-1 plasmid and linearized with SmaI (Tremblay et al., 1999). The probe was synthesized using a Promega riboprobe kit (Fisher Scientific, St-Laurent, QC), [^35^S]-UTP (Perkin Elmer, Woodbridge, ON) and polymerase T7 (Promega, Fisher Scientific, St-Laurent, QC). Labelled brain slices were placed against Kodak Biomax MR X-ray film (VWR, Town of Mount-Royal, QC) for 4 days.

### Quantification of c-fos mRNA

An experimenter blind to experimental condition translated optical gray densities from autoradiographs into microcuries (μCi) per gram of tissue using a ^14^C standard curve (ARC-146A, American Radiolabeled Chemicals, St-Louis, MI). Quantification was done using Image J software (NIH, Bethesda, MD). Background values were obtained from the rhinal fissure (+ 4.7 to 3.0 mm relative to Bregma) or the corpus callosum (+ 2.6 to 0.0 mm relative to Bregma) of each section. Background values were then subtracted from analysis. *C-fos* mRNA expression was measured in the orbitofrontal cortex [ventrolateral (VLO) and lateral (LO)], medial prefrontal cortex [prelimbic (PrL) and infralimbic (IL) cortex], cingulate cortex area 1 (Cg1), frontal cortex area 2 (FR2), caudate-putamen [(CPu; dorsomedial (DM), dorsolateral (DL), ventromedial (VM) and ventrolateral (VL) quadrants] and nucleus accumbens core (NacC) and shell (NacSh). Brain regions were identified according to (Paxinos and Watson, 1986). For each brain region, mRNA levels were averaged over 2-5 sections/rat.

### Statistical analysis

Two-way repeated-measures ANOVA was used to assess group differences in cocaine (or saline) intake, inter-infusion interval, lever presses and cumulative cocaine intake (Group x Session; the latter as a within-subjects variable). Two-way repeated-measures ANOVA was also used to assess group differences in number of lever presses and session length during PR testing (Group x Dose; the latter as a within-subjects variable). One-way ANOVA followed by Tukey’s multiple comparisons’ test or Kruskal-Wallis H test followed by Dunn’s multiple comparisons’ test were used to assess group differences in number of infusions, session length and locomotor activity on the last self-administration session before brain extraction. One-way ANOVA followed by Tukey’s multiple comparison test was used to analyse group differences in *c-fos* mRNA levels. Pearson’s r^2^ coefficients (two-tailed) were computed to assess relationships in c-*fos* mRNA expression between different brain regions.

## RESULTS

### LgA-90s rats take more cocaine than IntA-5s rats do, but the latter take cocaine at a faster pace

During the acquisition of cocaine self-administration behaviour prior to LgA-90s and IntA-5s sessions, projected LgA-90s rats completed more training days than IntA-5s rats did. This is because projected LgA-90s had to learn to self-administer cocaine infusions delivered over 5 s, with the timeout period gradually increasing from 20 to 85 s between sessions (data not shown; average number of days to acquisition; with 20-s timeout; 6 days ± 1 SEM; with 45-s timeout; 2 days; with 65-s timeout; 2 days; with 85-s timeout; 2 days). In contrast, projected IntA-5s rats only had to learn to self-administer cocaine infusions delivered over 5 s, with a 20-s timeout period (data not shown; average number of days to acquisition; 4 ± 1 SEM). As Figure 1 shows, after acquisition, rats from each projected group received three, 1-h cocaine self-administration sessions. Average cumulative number of self-administered infusions during all acquisition sessions, including the three, 1-h cocaine sessions was similar between the two cocaine groups (data not shown; projected IntA-5s rats, 309 ± 107 SEM; projected LgA-90s rats 233 ± 30 SEM; two-tailed *t*-test; *t*_(10)_ = 0.69, *p* = 0.51). This indicates that prior to starting IntA-5s and LgA-90s sessions, the rats assigned to these groups had experienced similar amounts of both cocaine and lever/cocaine/cue pairings.

After acquisition, the rats started IntA-5s or LgA-90s sessions. As **Figure 2A** illustrates, IntA would produce peaks and troughs in brain cocaine concentrations, while LgA would produce both higher and steadier brain drug concentrations (also see Allain et al., 2018; Zimmer et al., 2012). During self-administration sessions, IntA-5s rats were limited to 2 infusions/6-min cocaine period, providing a maximum of 24 cocaine infusions per 6-h session, as indicated by the dotted line in **Figure 2B**. In **Figures 2B-E,** the break between sessions 7 and 8 represents the four PR tests given between these sessions (data from these PR tests are reported under ‘*IntA-5s rats respond more for cocaine under a progressive ratio schedule of reinforcement than LgA-90s rats do’*). Rats self-administering saline during IntA or LgA sessions took similar numbers of injections and pressed an equivalent number of times on the active lever during their 6-h sessions (**Figures 2B** and **2D** respectively; all *P*’s > 0.05). Thus, we pooled these rats into a single control group (Sal-Ctrl) for subsequent analyses.

**Figure 2.**
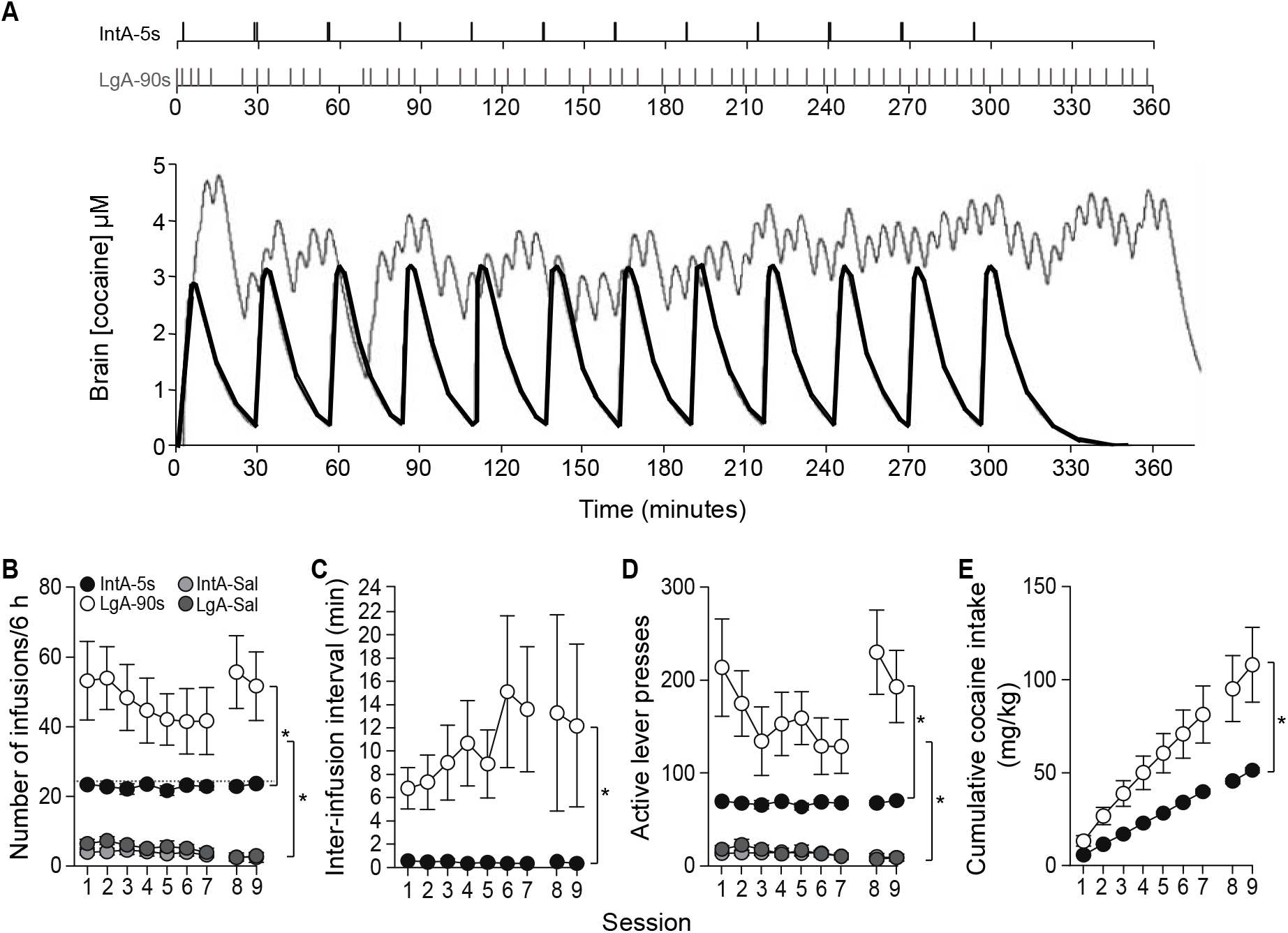
LgA-90s rats self-administered significantly more cocaine than IntA-5s rats did. IntA-5s rats were limited to 2 infusions (infused over 5 s) per 6-minute cocaine-available period. Each cocaine-available period was followed by a 26-minute no cocaine-available period (during which levers were retracted), for 12 cycles/session. LgA-90s rats had unlimited access to slower cocaine infusions (delivered over 90 s) during each 6-h session. (A) pattern of cocaine intake (top) and estimated brain cocaine concentrations (bottom) on the 7th day of IntA or LgA, from a representative rat from each group. (B) LgA-90s rats took more cocaine than IntA-5s rats did. (C) IntA-5s rats took cocaine at shorter inter-infusion intervals than LgA-90s rats did. (D) LgA-90s rats pressed more on the active lever than IntA-5s rats did. (E) cumulative cocaine intake was greatest in LgA-90s rats. All values are mean ± SEM. n = 7-21 rats/group. h, hours. IntA, intermittent access. LgA, long access. min, minutes. s, seconds. Sal, saline. *p < 0.05 vs. Saline or IntA-5s groups.

LgA-90s rats took significantly more cocaine than IntA-5s rats did (**Figure 2B**; Main effect of Group; F(1,17) = 13.99, *p* = 0.0016). Both cocaine-taking groups also self-administered more infusions than the saline groups did (Main effect of Group; IntA-5s vs. IntA-Sal, F(1,22) = 323.4; vs. LgA-Sal, F(1,19) = 254.3; LgA-90s vs. IntA-Sal, F(1,17) = 43.15; vs. LgA-Sal, F(1,14) = 29.99; all *P*’s < 0.0001).

IntA-5s rats took cocaine at a faster pace than LgA-90s rats did. **Figure 2C** shows the inter-infusion interval, calculated as the time elapsed between the end of one injection and the beginning of the next. For IntA-5s rats, the inter-infusion interval was computed during the 6-min cocaine periods. During these periods, IntA-5s rats took one injection every 26 s on average (± 5 s SEM). This was significantly shorter than the LgA-90s rats (11 min ± 4 min SEM; Main effect of Group; F(1,17) = 11.22, *p* = 0.004). The self-imposed inter-infusion interval in the LgA-90s rats was also significantly longer than the 90-s infusion length. This suggests that the 90-s infusion length did not artificially constrain cocaine self-administration behaviour in these rats.

Both cocaine groups pressed more on the active lever than saline control rats did (**Figure 2D**; Main effect of group; IntA-5s vs. IntA-Sal, F(1,22) = 286.5; vs. LgA-Sal, F(1,19) = 211.8; LgA-90s vs. IntA-Sal, F(1,17) = 43.91; vs. LgA-Sal, F(1,14) = 31.61; all *P*’s < 0.0001). LgA-90s rats also pressed more on the active lever than IntA-5s rats did (**Figure 2D**; F(1,17) = 18.46, *p* = 0.0005). LgA-90s rats pressed more on the inactive lever than all other groups did (LgA-90s vs. IntA-Sal, F(1,17) = 10.76; vs. LgA-Sal, F(1,14) = 7.27; vs. IntA-5s, F(1,17) = 10.36, all *P*’s < 0.02; data not shown).

Cumulative intake (number of injections taken × 0.25 mg/kg) was two times greater in LgA-90s rats than it was in IntA-5s rats (**Figure 2E**; Main effect of Group; F(1,17) = 14.83, *p* = 0.001). No other comparisons were statistically significant. Thus, compared to LgA-90s rats, IntA-5s rats took much less cocaine but at a significantly faster rate.

### IntA-5s rats respond more for cocaine under a progressive ratio schedule of reinforcement than LgA-90s rats do

Under a PR schedule of reinforcement, IntA-5s rats responded more for cocaine than LgA-90s rats did (**Figure 3A**; Dose x Group interaction effect; IntA-5s vs. LgA-90s: F(2,34) = 3.87, *p* = 0.03), in particular at the highest cocaine dose (*p* = 0.01). During these tests, Sal-Ctrl rats were self-administering saline, and IntA-5s and LgA-90s rats pressed more on the active lever than Sal-Ctrl rats did (**Figure 3A**; Main effect of Group; F(2,37) = 22.49; IntA-5s vs. Sal-Ctrl, F(1,31) = 42.55; LgA-90s vs. Sal-Ctrl, F(1,26) = 42.17, all *P*’s < 0.0001). In the cocaine groups, lever-pressing also increased with cocaine dose (Main effect of Dose; (2,74) = 11.03, p < 0.0001; IntA-5s vs. Sal-Ctrl, F(2,62) = 16.11, p < 0.0001; LgA-90s vs Sal-Ctrl, F(2,52) = 5.26, p = 0.008).

**Figure 3.**
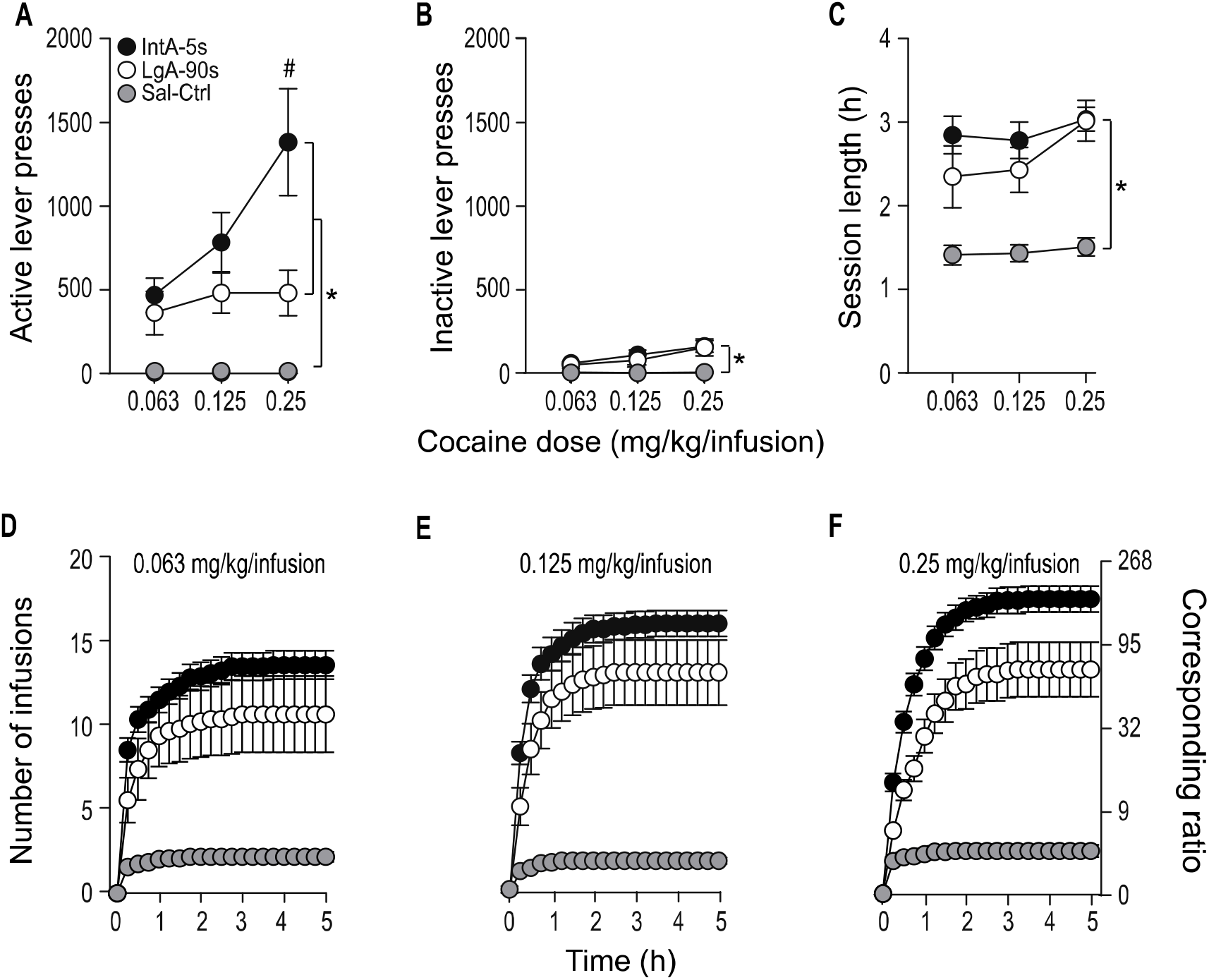
When tested under a progressive ratio schedule of reinforcement, IntA-5s rats responded more for cocaine than LgA-90s rats did. During progressive ratio sessions, IntA-5s and LgA-90s rats self-administered intravenous cocaine, and Sal-Ctrl rats self-administered saline. All infusions were delivered over 10 s. (A) Responding on the active lever was greatest in IntA-5s rats. (B) Both IntA-5s and LgA-90s rats responded more on the inactive lever than Sal-Ctrl rats did. (C) Sessions lasted longer in IntA-5s and LgA-90s rats than they did in Sal-Ctrl rats. (D-F) IntA-5s responded more for cocaine than LgA-90s did, throughout each progressive ratio test session, and across a range of cocaine doses, as indicated by a greater cumulative number of infusions taken over 15-minute bins during each test session. All values are mean ± SEM. n = 7-21 rats/group. h, hours. IntA, intermittent access. LgA, long access. s, seconds. Sal-Ctrl, saline control animals. *p < 0.05 vs. Sal-Ctrl group. #p < 0.05 vs. IntA-5s group

IntA-5s and LgA-90s animals did not differ in the number of inactive lever presses during PR sessions, and both groups pressed more on the inactive lever than Sal-Ctrl rats did (**Figure 3B**; Main effect of Group: F(2,37) = 14.15, IntA-5s vs. Sal-Ctrl, F(1,31) = 30.37; ContA-90 s vs. Sal-Ctrl, F(1,26) = 24.56; all *P*’s < 0.0001) and pressing on the inactive lever also increased with cocaine dose (Main effect of Dose; F(2,74) = 35.78; *p* < 0.05). PR sessions lasted longer in the IntA-5s and LgA-90s rats than they did in Sal-Ctrl rats (**Figure 3C**; F(2,37) = 42.72; all P’s < 0.002). PR sessions also lasted longer at higher cocaine doses (Main effect of dose; F(2,34) = 3.87, *p* = 0.03). No other comparisons were statistically significant. **Figures 3D-F** show cumulative number of drug infusions taken during the 5-h progressive-ratio sessions, as a function of time and cocaine dose. Visual inspection of **Figures 3D-F** suggests that, in particular at the highest dose tested, the IntA-5s rats earned more cocaine than the LgA-90s rats, at all time points during the PR session. Thus, IntA-5s rats had taken significantly less cocaine in the past than LgA-90s rats had, but IntA-5s rats later showed greater incentive motivation for the drug.

### Self-administration behaviour and locomotor activity on the last session before brain extraction

On the final self-administration session before brain extraction, rats could self-administer a maximum of 10 infusions, each delivered over 10 s. The Sal-Ctrl rats self-administered saline. The session ended once 10 infusions were taken or after a maximum of 1 h. **Figure 4A** shows that IntA-5s and LgA-90s rats took a similar number of cocaine infusions (χ^2^_(2)_ = 23.93, *p* > 0.9999), and both groups self-administered more infusions than Sal-Ctrl rats did (χ^2^_(2)_ = 23.93, *p* < 0.0001; IntA-5s vs. Sal-Ctrl, *p* < 0.0001; LgA-90s vs. Sal-Ctrl, *p* = 0.004). The self-administration session was shorter in IntA-5s rats than it was in Sal-Ctrl rats (**Figure 4B**; (χ^2^_(2)_ = 22.31, *p* < 0.0001; IntA-5s vs. Sal-Ctrl: *p* <0.0001; LgA-90s vs. Sal-Ctrl *p* = 0.12). Both session length (*p* = 0.29) and inter-infusion interval (data not shown; Unpaired *t*-test; *t* _(7,68)_ = 1.98, *p* = 0.08) were similar in the IntA-5s and LgA-90s rats. The two groups also showed similar levels of cocaine-induced locomotion during this final self-administration session, and both showed greater locomotion than the Sal-Ctrl rats did (**Figure 4C**; χ^2^_(2)_ = 27.21, *p* < 0.0001; IntA-5s vs. LgA-90s, *p* > 0.9999, IntA-5s vs. Sal-Ctrl, *p* < 0.0001 and LgA-90s vs. Sal-Ctrl, *p* = 0.0008).

**Figure 4.**
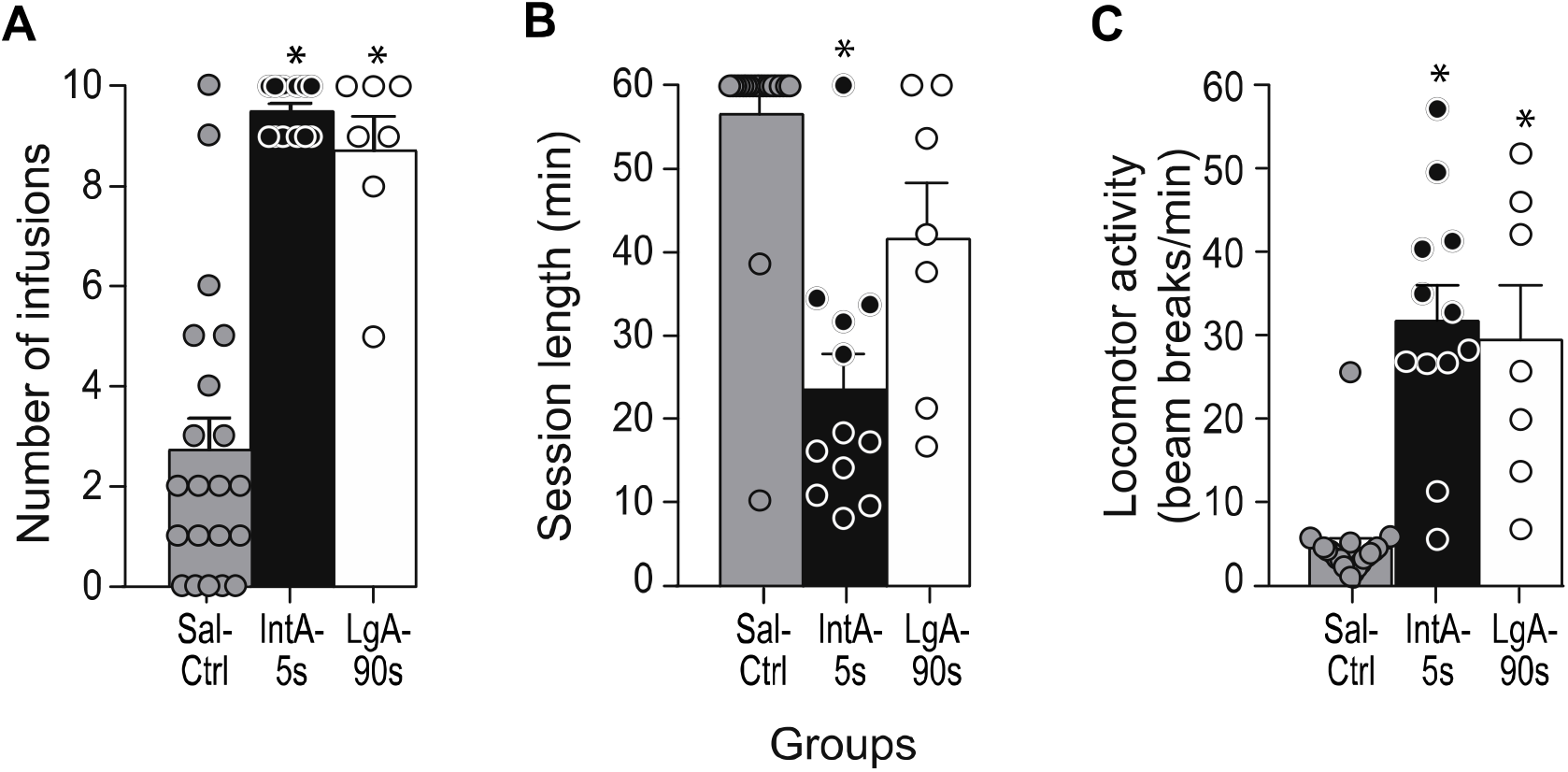
On the final self-administration session prior to brain extraction, IntA-5s rats and LgA-90s rats consumed similar amounts of cocaine and showed similar levels of drug-induced locomotion. During the final self-administration session prior to brain extraction, all rats were limited to a maximum of 10 intravenous cocaine infusions (0.25 mg/kg/infusion; or saline for control rats) delivered over 10 s. Sessions ended once the infusion criterion was met or after 1 hour. The two cocaine-taking groups (A) took similar amounts of cocaine, (B) had similar session lengths and (C) showed similar levels of cocaine-evoked locomotor activity. All values are mean ± SEM. n = 7-21 rats/ group. IntA, intermittent access. LgA, long access. min, minutes. s, seconds. Sal-Ctrl, saline control animals. *p < 0.05 vs. Sal-Ctrl group.

Thus, during the last self-administration session before brain extraction, IntA-5s rats and LgA-90s rats took similar amounts of cocaine and showed similar levels of drug-induced locomotion. A subset of the rats in **Figure 4** were used for *in situ* hybridization (IntA-5s and LgA-90s, n = 6/group; Sal-Ctrl, n = 10). During the last self-administration session before brain extraction, this subset of IntA-5s rats and LgA-90s rats had similar levels of cocaine intake (**Figure S1A**; χ^2^_(2)_ = 16.03, *p* > 0.9999), session length (**Figure S1B**; F(2,19) = 13.15, p = 0.18), and drug-induced locomotion (**Figure S1C**; χ^2^_(2)_ = 15.65, *p* > 0.9999).

### *Compared to LgA-90s rats, IntA-5s rats show greater cocaine-induced c*-fos *mRNA expression in the frontal cortex and the striatum*

In the orbitofrontal, medial prefrontal, cingulate and frontal cortices, cocaine self-administration increased *c-fos* mRNA expression compared to saline self-administration (**Figure 5;** all *P*’s < 0.05). The mode of drug intake in the past also determined cocaine-induced *c-fos* mRNA levels. IntA-5s rats had more *c-fos* mRNA than LgA-90s rats did in the ventrolateral and lateral orbitofrontal cortices and in the prelimbic cortex (**Figures 5B-D**; VLO: F(2,107) = 57.5; LO: F(2,63) = 32.57; PrL: F(2,85) = 37.88; all *P*’s < 0.0001). There were no differences between these groups in the infralimbic, cingulate or frontal area 2 cortices (**Figures 5E-G**; all *P*’s > 0.05).

**Figure 5.**
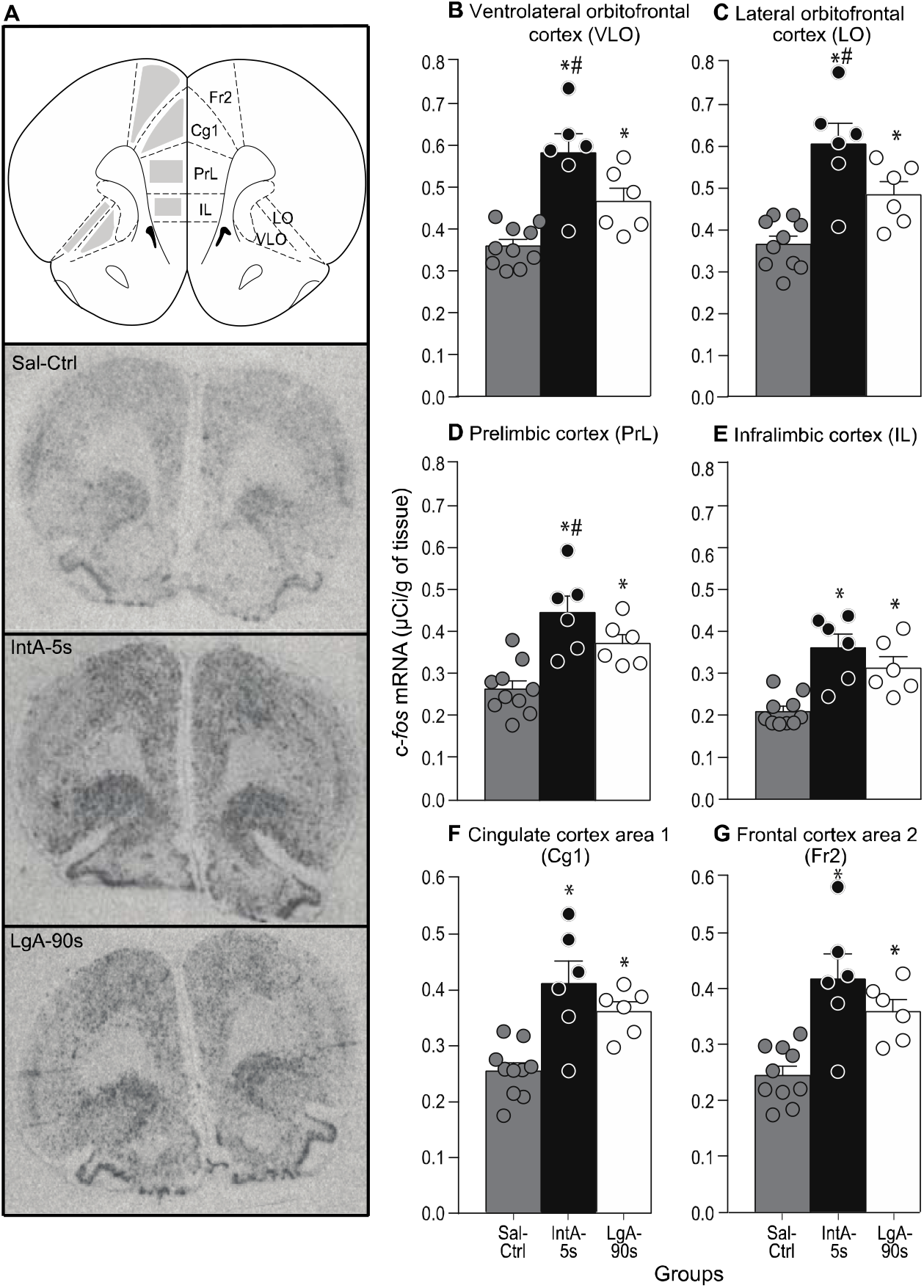
IntA-5s rats have more cocaine-induced c-*fos* mRNA expression in the orbitofrontal and prelimbic cortices than LgA-90s rats do. (**A**) The regions quantified and autoradiographs for c*-fos* mRNA expression in a sample rat from each group. Compared to LgA-90s rats, IntA-5s rats showed higher levels of cocaine-evoked c*-fos* mRNA density in (**B**) the ventrolateral orbitofrontal cortex, (**C**) the lateral orbitofrontal cortex and (**D**) the prelimbic cortex. Compared to saline control rats, both IntA-5s and LgA-90s rats showed more c*-fos* mRNA expression in (**E**) the infralimbic cortex, (**F**) the cingulate cortex, area 1, and (**G**) the frontal cortex area 2, but there were no differences between the cocaine-taking groups in these brain regions. All values are mean ± SEM. *n* = 6-10 rats/group. *Sal-Ctrl*, saline control animals. *IntA*, intermittent access. *LgA*, long access. *s*, seconds. *VLO*, ventrolateral orbitofrontal cortex. *LO*, lateral orbitofrontal cortex. *PrL*, prelimbic cortex. *IL*, infralimbic cortex. *Cg1*, cingulate cortex area 1. *Fr2*, frontal cortex area 2. \**p* < 0.05 vs. Sal-Ctrl group. ^#^*p* < 0.05 vs. LgA-90s group.

Cocaine self-administration increased *c-fos* mRNA expression in the caudate putamen (**Figures 6B-E**) and in the nucleus accumbens (**Figures 6F-G**) compared to saline self-administration (all *P’*s < 0.05). Compared to LgA-90s rats, IntA-5s rats also had increased *c-fos* mRNA levels in all quadrants of the caudate-putamen (**Figures 6B-E**; DM: F(2.85) = 35.14; DL: F(2.85) = 27.66; VM: F(2.85) = 41.38; VL: 31.78; all *P*’s < 0.0001). There were no differences between these groups in the nucleus accumbens core or shell (**Figures 6F-G**; all *P*’s > 0.05).

**Figure 6.**
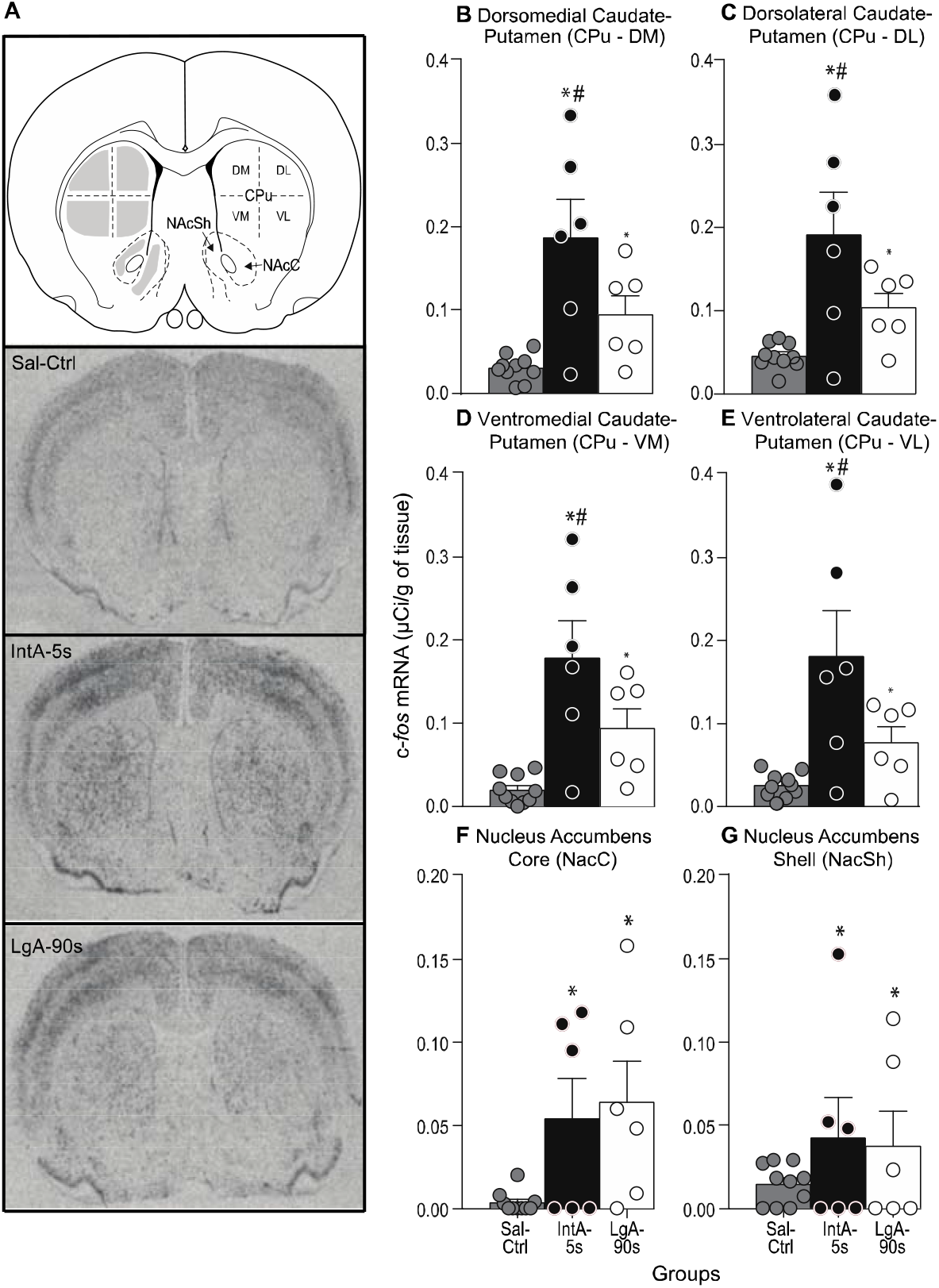
IntA-5s rats show more cocaine-induced c-*fos* mRNA expression in the caudate-putamen than LgA-90s rats do. (**A**) The regions quantified and autoradiographs for c*-fos* mRNA expression in a sample rat from each group. Compared to LgA-90s rats, IntA-5s rats showed higher levels of cocaine-evoked c*-fos* mRNA density in (**B**) the dorsomedial caudate-putamen, (**C**) the dorsolateral caudate-putamen, (**D**) the ventromedial caudate-putamen, and (**E**) the ventrolateral caudate-putamen. Compared to saline control rats, both IntA-5s and LgA-90s rats showed more c*-fos* mRNA expression in (**F**) the nucleus accumbens core and (**G**) the nucleus accumbens shell, but there were no differences between the cocaine-taking groups in these brain regions. All values are mean ± SEM. *n* = 6-10 rats/group. *Sal-Ctrl*, saline control animals. *IntA*, intermittent access. *LgA*, long access. *s*, seconds. *CPu*, caudate putamen. *DM*, dorsomedial quadrant. *DL*, dorsolateral quadrant. *VM*, ventromedial quadrant. *VL*, ventrolateral quadrant. *NacC*, nucleus accumbens core. *NacSh*, nucleus accumbens shell. \**p* < 0.05 vs. Sal-Ctrl group. ^#^*p* < 0.05 vs. LgA-90s group.

### *C*-fos *mRNA levels in the frontal cortex and the dorsal striatum are positively correlated only in IntA-5s rats*

Corticostriatal afferents are a massive source of glutamate in the striatum (Berendse et al., 1992; Carter, 1982; Gerfen, 1992; McGeorge and Faull, 1989) and psychostimulant-induced immediate early gene expression in the striatum depends on glutamatergic inputs from the cortex (Cenci and Björklund, 1993; Ferguson and Robinson, 2004; Hess et al., 2003). The prelimbic cortex sends excitatory inputs to the *dorsomedial* caudate-putamen (Berendse et al., 1992; Mailly et al., 2013). In contrast, the ventrolateral and lateral areas of the orbitofrontal cortex send glutamatergic afferents preferentially to the *ventro/centrolateral* caudate-putamen (Berendse et al., 1992; Mailly et al., 2013; Schilman et al., 2008). **Figure 7** shows correlational analyses between c-*fos* mRNA levels in these cortical regions and the caudate-putamen. In IntA-5s rats only, more cocaine-induced c-*fos* mRNA in the ventrolateral and lateral orbitofrontal cortex significantly predicted more cocaine-induced immediate early gene expression in the ventrolateral caudate-putamen (**Figure 7A**, *r*^*2*^ = 0.85, *p* = 0.009). Similarly, in the IntA-5s group only, more cocaine-induced c-*fos* mRNA in the prelimbic cortex significantly predicted more cocaine-induced immediate early gene expression in the dorsomedial caudate-putamen (**Figure 7D**, *r*^*2*^ = 0.94; *p* = 0.001). Thus, cocaine-induced gene regulation in frontal cortical regions and in the caudate-putamen was significantly correlated *only* in rats with a history of taking rapid and intermittent cocaine injections.

**Figure 7.**
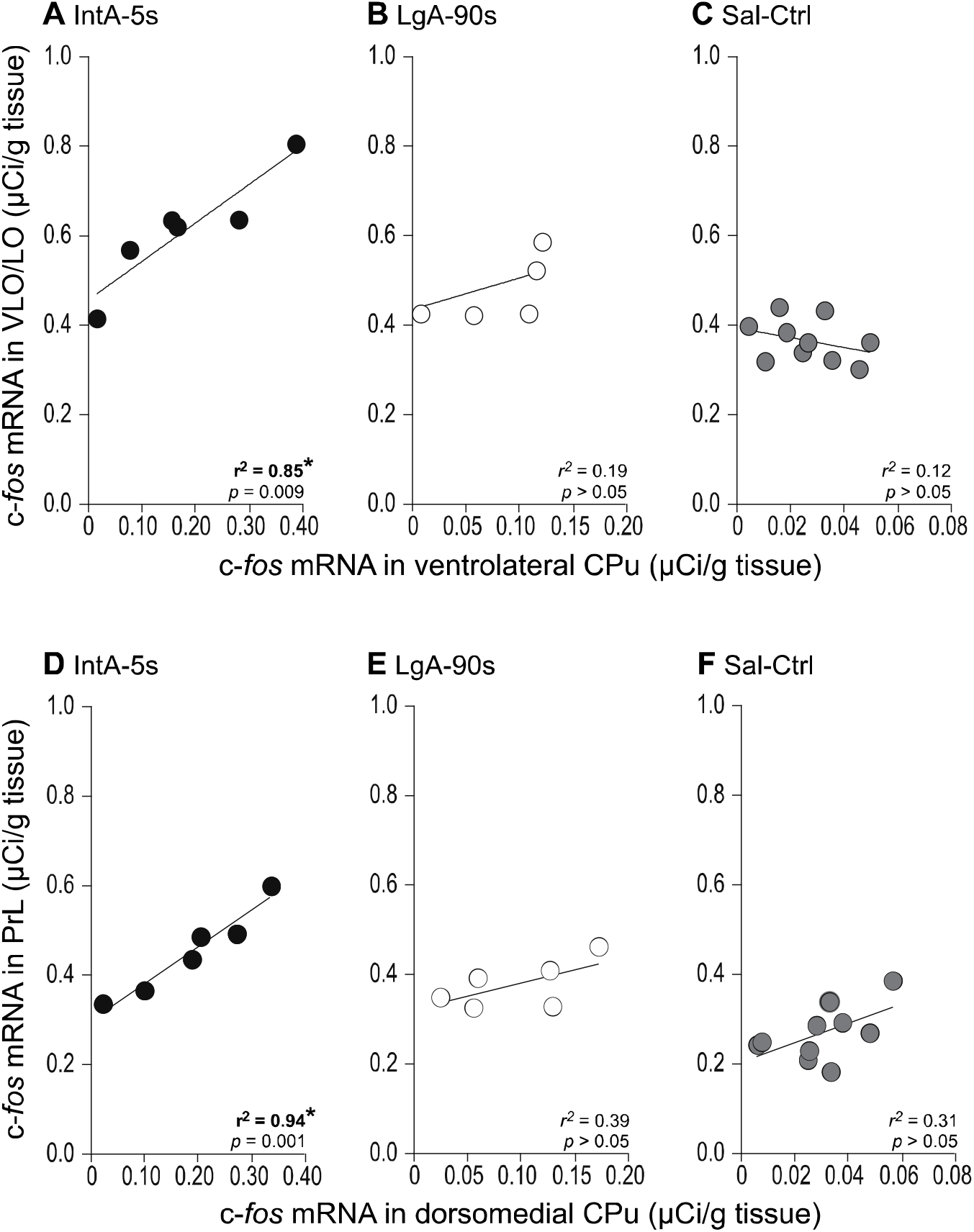
Only IntA-5s rats show significant positive correlations between cocaine-induced c-*fos* mRNA levels in frontal cortical regions and in the caudate-putamen. C-*fos* mRNA expression induced by cocaine self-administration was significantly correlated between the orbitofrontal cortex and the ventrolateral caudate-putamen in (**A**) IntA-5s rats, but not in (**B**) LgA-90s or (**C**) Sal-Ctrl rats. Similarly, c-*fos* mRNA expression induced by cocaine self-administration was significantly correlated between the prelimbic cortex and the dorsomedial caudate-putamen in (**D**) IntA-5s rats, but not in (**E**) LgA-90s or (**F**) Sal-Ctrl rats. *Sal-Ctrl*, saline control animals. *IntA*, intermittent access. *LgA*, long access. *s*, seconds. CPu, caudate-putamen. *VLO*, ventrolateral orbitofrontal cortex. *LO*, lateral orbitofrontal cortex. *PrL*, prelimbic cortex.

## DISCUSSION

Here we compared outcome in rats given intermittent access to rapid cocaine injections (IntA-5s) and in rats given continuous access to slower injections (LgA-90s). We report three main findings. First, IntA-5s rats took two times less cocaine than LgA-90s rats did, but IntA-5s rats showed greater incentive motivation for the drug, as measured by responding for cocaine under a PR schedule of reinforcement. Second, cocaine self-administration increased c-*fos* mRNA levels in corticostriatal regions of both LgA-90s and IntA-5s rats, but IntA-5s rats had comparatively greater gene regulation in the OFC, prelimbic cortex and in the caudate-putamen. Third, cocaine-induced c-*fos* mRNA expression in frontal cortical regions and in the caudate-putamen were significantly correlated *only* in IntA-5s rats. These findings demonstrate that the mode of cocaine intake influences both incentive motivation for the drug and cocaine-induced gene regulation in the brain, and that greater motivation to take cocaine is potentially linked to increased drug-induced recruitment of corticostriatal regions.

### LgA-90s rats self-administer more cocaine but IntA-5s rats show more incentive motivation for the drug

The LgA-90s rats consumed significantly more cocaine than IntA-5s rats did, but they did not escalate their cocaine intake over sessions. The lack of escalation is similar to other LgA studies conducted in males and using lower cocaine doses (0.25-0.6 mg/kg/infusion; Ferrario and Robinson, 2007; Kippin et al., 2006; Mantsch et al., 2004; Minogianis et al., 2013). Our IntA-5s rats also did not escalate their intake over sessions. IntA experience can produce escalation of intake (Allain et al., 2018; Allain and Samaha, 2018; James et al., 2018; Kawa et al., 2016; Kawa et al., 2018; Pitchers et al., 2017a; Pitchers et al., 2017b; Singer et al., 2018). However, here we limited cocaine intake during each IntA session, and this precludes escalation (also see Allain et al., 2018). Still, even though IntA-5s rats had previously taken half the amount of cocaine as LgA-90s had, IntA-5s rats subsequently worked harder to obtain cocaine when physical cost was high (i.e., under a PR schedule of reinforcement). In this regard, it is worth noting that using behavioural economics measures of cocaine demand, it was shown that LgA can produce a persistent increase in the preferred level of cocaine consumption when cost is low, but only a transient increase in the motivation to obtain cocaine (James et al., 2018). In contrast, the same study showed that IntA does not change preferred level of intake, but it produces a very persistent increase in the motivation for cocaine (up to 50 days after IntA experience; James et al., 2018). Thus, high levels of cocaine exposure in the past do not necessarily produce greater cocaine wanting in the future, and intermittent intake of rapid drug injections is particularly effective in enhancing incentive motivation for cocaine (also see Allain et al., 2018; Zimmer et al., 2012). This is consistent with a growing literature showing that how fast and how often cocaine is consumed are more important than how much drug is taken in promoting addiction-like symptoms [(Allain et al., 2018; Allain et al., 2017; Bouayad-Gervais et al., 2014; James et al., 2018; Kawa et al., 2016; Liu et al., 2005; Minogianis et al., 2013; Singer et al., 2018; Wakabayashi et al., 2010; Zimmer et al., 2012), reviewed in (Allain et al., 2015; Kawa et al., 2019)].

Intermittent, rather than continuous cocaine intake might also be more clinically relevant. It is unlikely that cocaine users maintain continuously high brain concentrations of drug when consuming cocaine, as achieved by LgA procedures. Instead, cocaine use in humans is defined by intermittency, both between and within bouts of consumption (Beveridge et al., 2012; Cohen and Sas, 1994; Simon et al., 2001). Our present findings and this literature suggest that IntA procedures might most useful to study the changes in brain, psychological function and behaviour involved in the development of cocaine addiction.

### *The mode of past cocaine use produces both qualitative and quantitative differences in cocaine-induced c-*fos *mRNA expression*

In almost every brain region studied, IntA-5s rats showed more cocaine-evoked c-*fos* mRNA levels than LgA-90s rats did. This cannot be explained by any acute differences in the mode of cocaine intake, achieved dose or motor activity immediately prior to c-*fos* mRNA quantification. Indeed, on the last self-administration session before brain extraction, the two groups had access to cocaine under the same conditions (continuous access to i.v. cocaine infusions injected over 10 s), they took similar amounts of cocaine and they also showed similar levels of cocaine-induced psychomotor activity. Instead, the differences in cocaine-induced c-*fos* expression are likely due to the pattern of past cocaine intake. LgA-90s rats achieved high and sustained brain cocaine concentrations during each session, and they also had greater cumulative drug exposure than IntA-5s rats. IntA-5s rats achieved repeated ‘spikes’ in brain cocaine concentrations, and also took comparatively little drug. Repeated exposure to high and steady cocaine concentrations in the LgA-90s rats could have blunted the c-*fos* mRNA response to subsequent cocaine. Determining this conclusively would require assessing cocaine-induced c-*fos* mRNA expression in previously cocaine-naïve rats. Previous work does show that compared to cocaine-naïve rats, rats that have repeatedly self-administered i.v. cocaine (Daunais et al., 1993; Daunais et al., 1995; Ennulat et al., 1994) or that have received repeated intraperitoneal cocaine injections [(Daunais and McGinty, 1994; Hope et al., 1992; Steiner and Gerfen, 1993), (reviewed in Hammer et al., 1997)] show tolerance to cocaine-induced c-*fos* mRNA expression. It is possible then, that the LgA-90s condition more effectively promotes this tolerance compared to the IntA-5s condition. This is especially likely when one considers the very different effects LgA and IntA cocaine experience produce within the dopamine system. Psychostimulant drugs increase immediate early gene expression in part by enhancing dopamine neurotransmission (Ferguson et al., 2003; Graybiel et al., 1990; Ruskin and Marshall, 1994; Young et al., 1991), and LgA cocaine experience promotes tolerance to cocaine’s dopamine-elevating effects, while IntA cocaine experience promotes sensitization (Allain et al., 2020; Calipari et al., 2014; Calipari et al., 2013; Ferris et al., 2011; Kawa et al., 2018; Siciliano et al., 2018). A possibility here is that the c-*fos* mRNA response we measured reflects the neural response not only to cocaine intake, but to the cocaine-paired context/cues as well. Indeed, all rats were exposed to the self-administration chamber prior to brain extraction. The self-administration context could have contributed to the c-*fos* response especially in IntA-5s rats, because these rats were sacrificed sooner after being introduced to the self-administration chamber than LgA-90s rats were. That is, the final self-administration session prior to sacrifice tended to be shorter in IntA-5s versus LgA-90s rats (Figure S1B). Future work could directly compare context/cue-induced c-*fos* mRNA levels in these groups. This is an especially interesting avenue for future research, because compared to LgA rats, IntA rats attribute more incentive motivational properties to cocaine-paired cues (Nicolas et al., 2019).

The group differences in cocaine-induced c-*fos* mRNA expression could also involve the intake of rapid versus slower cocaine infusions specifically, independent of any differences in the intermittency of intake. Indeed, the present results do not allow us to parse out the influence of the pattern (intermittency) of past cocaine intake versus the speed of past cocaine intake, and future work is needed to address this. As such work unfolds, it is known that at least when cocaine is administered acutely, increasing the speed of i.v. infusion promotes drug-induced c-*fos* mRNA expression in the medial prefrontal and orbitofrontal cortices (Samaha et al., 2004) and in the striatum (Ferrario et al., 2008; Samaha et al., 2004). We highlight, however, that here LgA-90s rats and IntA-5s rats self-administered cocaine infusions delivered at a common speed immediately prior to c-*fos* mRNA processing. Thus, if group differences in cocaine-induced c-*fos* mRNA involve the speed of drug delivery, this is likely because the intake of rapid versus slower cocaine infusions *in the past* determine the neurobiological impact of subsequent exposure to the drug. In support, the intake of rapid versus slower cocaine infusions evokes different long-term adaptations in both dopamine D2 receptor number and function in the striatum (Minogianis et al., 2013). Whatever the precise mechanisms involved, a unique finding here is that in rats that show enhanced incentive motivation to take cocaine (IntA-5s rats), there is augmented cocaine-induced gene regulation in the frontal cortex and striatum. This suggests that potentiated cocaine-induced recruitment of these brain regions could mediate the expression of the enhanced cocaine wanting that defines addiction.

The mode of cocaine intake also produced *qualitative* differences in cocaine-evoked c-*fos* mRNA expression. There was a significant positive correlation between cocaine-induced c-*fos* mRNA expression in the orbitofrontal/prelimbic cortices and that in the caudate-putamen *only* in IntA-5s rats. This pattern of apparent connectivity unique to IntA-5s rats could reflect the extent to which specific neural circuits are recruited when cocaine has increased incentive motivational value. Three lines of evidence support this idea. First, the OFC encodes the motivational value of rewards (Tremblay and Schultz, 1999; Wallis and Miller, 2003), and sends this information to the striatum to guide responding to rewards (Pennartz et al., 2000; Schultz et al., 2000). It follows then that if rats assign increased motivational value to cocaine, this should be reflected by enhanced cocaine-induced recruitment of the OFC and striatum. Second, substantial monosynaptic glutamatergic inputs connect the OFC to the caudate-putamen (Berendse et al., 1992; Carter, 1982; Gerfen, 1992; McGeorge and Faull, 1989; Schilman et al., 2008), and glutamate from corticostriatal afferents mediates drug-induced c-*fos* mRNA in the caudate-putamen, particularly when drugs are experienced under conditions that promote sensitization-related changes in brain and behaviour (Ferguson et al., 2003; Ferguson and Robinson, 2004; Snyder-Keller, 1991). Third, recent findings show that in IntA-5s rats, transiently disconnecting the OFC and the caudate-putamen with GABA receptor agonists suppresses incentive motivation to take cocaine (Minogianis et al., 2019), as measured by responding for the drug under a PR schedule of reinforcement. Together, the findings suggest that the roles of the OFC, caudate-putamen and their connections in incentive motivation to take cocaine should be examined further, using circuit-specific chemogenetic or optogenetic manipulations for example. Such work would be an important complement to recent discoveries about the neurochemical and neuroanatomical underpinnings of the IntA phenotype. For example, a new study shows that IntA (but not LgA) cocaine experience persistently increases lateral hypothalamic orexin cell function in ways that promote addiction-like behavior, and that this can be reversed by knockdown of orexin neurons (James et al., 2018).

## CONCLUSIONS

Our findings demonstrate that rats given continuous access to sustained i.v. cocaine infusions take large amounts of the drug, while rats given intermittent access to rapid cocaine infusions show more incentive motivation to take cocaine, even though they have taken comparatively little drug overall. Moreover, self-administered cocaine engages the OFC, prelimbic cortex and caudate-putamen most markedly in rats that show enhanced incentive to obtain the drug, with apparently increased corticostriatal connectivity in these same animals. We conclude that enhanced cocaine-induced recruitment of corticostriatal circuits is potentially involved in the expression of the increased cocaine wanting that defines the addicted state.

## ACKNOWLEDGEMENTS AND FUNDING

We would like to thank Alice Servonnet and David Voyer for preparing the plasmid. This work was supported by grants from the Canadian Institutes of Health Research (Grant No. 157572) and the Canadian Foundation for Innovation (Grant No. 24326) awarded to ANS. ANS holds a salary award from the Fonds de la Recherche du Québec - Santé (Grant No. 28988). EAM holds a graduate fellowship from Fonds de la Recherche du Québec - Santé (Grant No. 29651).

## AUTHOR CONTRIBUTIONS

EAM and ANS designed the study. EAM performed all experiments. EAM and ANS analyzed the data and interpreted the findings. EAM and ANS drafted the manuscript. All authors critically reviewed content and approved final version for publication

---

**Supplementary figure 1.**
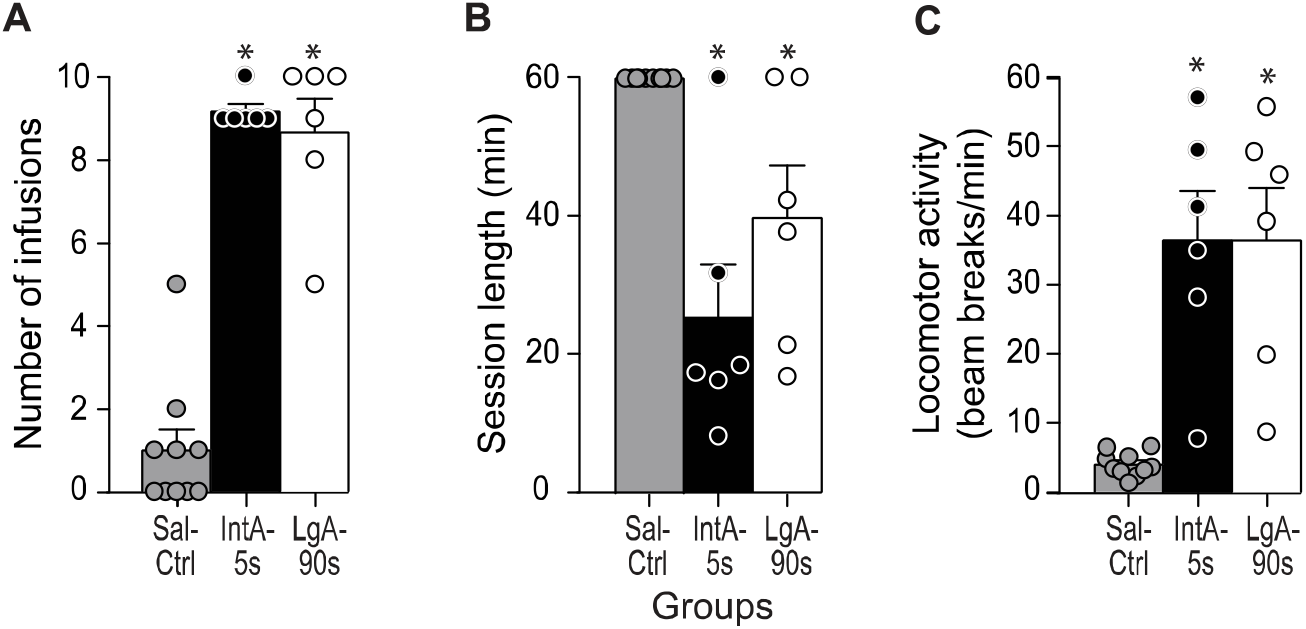
Self-administration data from the subsets of rats used to quantify c-*fos* mRNA expression. During the final self-administration session prior to brain extraction, all rats were limited to a maximum of 10 intravenous cocaine infusions (0.25 mg/kg/infusion; or saline for control rats delivered over 10 s. Sessions ended once the infusion criterion was attained or after 1 hour. The two cocaine-taking groups (**A**) took similar amounts of cocaine, (**B**) had similar sessio lengths and (**C**) showed similar levels of cocaine-evoked locomotor activity. All values are mean ± SEM. n = 6-10 rats/group. IntA, intermittent access. LgA, long access. min, minutes. s, seconds. Sal-Ctrl, s aline control rats. *p < 0.05 vs. Sal-Ctrl group.

